# Biological network topology features predict gene dependencies in cancer cell lines

**DOI:** 10.1101/751776

**Authors:** Graeme Benstead-Hume, Sarah K. Wooller, Samantha Dias, Lisa Woodbine, Anthony M. Carr, Frances M. G. Pearl

## Abstract

In this paper we explore computational approaches that enable us to identify genes that have become essential in individual cancer cell lines. Using recently published experimental cancer cell line gene essentiality data, human protein-protein interaction (PPI) network data and individual cell-line genomic alteration data we have built a range of machine learning classification models to predict cell line specific acquired essential genes. Genetic alterations found in each individual cell line were modelled by removing protein nodes to reflect loss of function mutations and changing the weights of edges in each PPI to reflect gain of function mutations and gene expression changes.

We found that PPI networks can be used to successfully classify human cell line specific acquired essential genes within individual cell lines and between cell lines, even across tissue types with AUC ROC scores of between 0.75 and 0.85. Our novel perturbed PPI network models further improved prediction power compared to the base PPI model and are shown to be more sensitive to genes on which the cell becomes dependent as a result of other changes. These improvements offer opportunities for personalised therapy with each individual’s cancer cell dependencies presenting a potential tailored drug target.

The overriding motivation for predicting cancer cell line specific acquired essential genes is to provide a low-cost approach to identifying personalised cancer drug targets without the cost of exhaustive loss of function screening.

## Introduction

An essential gene is one which is necessary for cellular survival and reproductive success. However, the exact set of essential genes is context specific depending on the cell type, genetic and epigenetic aberrations and the cell environment. The different definitions and measurements of essentiality often have considerable overlap but there are also large areas of disagreement (Eisenberg & Levanon, 2013; Bartha *et al*, 2018).

During the process of carcinogenesis, the pattern of essential genes changes as cells become addicted to oncogenes and tumour suppressor genes become inactivated (Weinstein, 2002; Luo *et al*, 2009). Identifying gene dependencies that result from carcinogenesis can provide opportunities for targeted treatments, as the inhibition of proteins which are essential in cancer cells but not in normal cells can lead to selective cell death (Workman *et al*, 2013). However, the heterozygous nature of cancer and the large number of genetic alterations in cancer cell lines prevents the exhaustive identification of these acquired essential proteins for all possible cell lines.

Several groups have used features derived from protein-protein interaction (PPI) networks to predict cancer genes (Li *et al*, 2009), and genetic interactions (Benstead-Hume *et al*, 2019). Furthermore there have been a number of successful attempts to predict common essential genes using biological network data in different contexts and in different organisms (for a review see Zhang et al. (Zhang *et al*, 2016)). These studies have used a range of different network data including protein-protein interaction (PPI) networks, transcriptional regulatory networks, gene co-expression networks, metabolic networks (MNs) and networks that integrate two or more of the above. Due to data availability these studies have generally focused on model organisms. For studies on *S. cerevisiae* (Chen & Xu, 2005; Saha & Heber, 2006; Acencio *et al*, 2009). For studies on *E. coli* see *(da Silva *et al*, 2008; Hwang *et al*, 2009)* and for studies on various bacteria see (Plaimas *et al*, 2010; Cheng *et al*, 2014; Lu *et al*, 2014). For the most part these studies employ similar methods where topology data is extracted from the biological networks. This topology data is subsequently used as a feature set to train machine learning models to identify essential genes. For example, Saha et al. (Saha & Heber, 2006) reported a ROC AUC of 82% using PPI network degree count and conservation score features to classify ∼2,200 essential genes in *S. cerevisiae* and Müller da Silva *et al*. (da Silva *et al*, 2008) who reported F1 scores of 83.4% for essential gene predictions and 79.7% for non-essential gene prediction in *E coli*. Similar predictions have not been reported for human cell lines.

Generally past studies have focused on a static version of the known PPI network with little modification for individual samples. Observations made by Roumeliotis et al. (Roumeliotis *et al*, 2017), that suggest the effect of genetic variations can be transmitted from directly affected proteins to distant gene products through protein interaction pathways, suggest that the inclusion genetic alterations may allow us to improve the traditional PPI network model.

Recently there have been significant experimental efforts to identify and catalogue cancer specific acquired essential genes, otherwise known as gene dependencies, experimentally. Amongst these efforts are a number of loss of function screens (Ngo *et al*, 2006) performed using both RNAi and CRISPR-Cas9 systems (Marcotte *et al*, 2016, 2012; Luo *et al*, 2008; Cheung *et al*, 2011; Aksoy *et al*, 2014; Aguirre *et al*, 2016). These screens investigate the changes in phenotype caused in cell lines by systematically knocking genes out one by one either through deletion or disruption. Knock-outs that result in significantly deleterious phenotypes signal that the respective gene may be essential in that cell line.

In response to reported off-target effects observed in loss of function screens, where genes other than the target are disrupted by certain RNAi (Jackson & Linsley, 2004; Birmingham *et al*, 2006; Buehler *et al*, 2012; Munoz *et al*, 2016; Aguirre *et al*, 2016), Tsherniak et al. (Tsherniak *et al*, 2017) building on previous work by Cowley et al. (Cowley *et al*, 2014), performed 285 genome scale systematic loss-of-function screens to identified cancer dependencies across a total of 501 human cancer cell lines covering 21 different tissue types. They found 6,476 genes that had a cancer dependency score of over 0.65 in at least one cell line. Of these 6,476 genes, 545 were dependencies in 20-50% of cell lines in at least one tissue-type. This suggested that these genes are commonly essential in cancer cells of that tissue type but non-essential in normal cells.

While identifying general essential genes or disease specific gene dependencies provides a better understanding of potential disease specific targets, loss of function screens are not readily available for the majority of individual cancer patients. Tools that could predict cell line specific gene dependencies from more readily available data such as mutations and gene expression may offer new opportunities for affordable tailored therapies (Charlton & Spicer, 2016; Benstead-Hume *et al*, 2017).

In this study we use recent cell line specific gene dependency data along with PPI networks data to build models able to identify novel cell line specific gene dependencies. To do this we model genetic alterations in specific cell lines by perturbing their respective PPI networks. We explore the viability of identifying cell line specific gene dependencies both within and between various human cancer cell lines using this perturbed PPI networks data. Finally, we introduce DependANT, a classifier trained to predict cell line specific gene dependencies using both generic and perturbed PPI networks data with the aim of providing a low cost approach to identifying personalised cancer drug targets without the cost of exhaustive loss of function screening.

## Results

### Data sets

DependANT classifies cell line specific gene dependencies via models built using protein-protein interaction (PPI) network and genetic alteration data. The PPI networks were sourced via STRING (von Mering *et al*, 2005) and the mutation and gene expression data used to perturb our networks, as well as the gene dependency scores used to label our training data, are publicly available from Tsherniak et al. via project Achilles (Tsherniak *et al*, 2017).

We selected all breast, kidney and pancreatic cancer cell lines that had sufficient gene dependency and genetic alteration data in the Tsherniak data (figure EV 1). These included 19 breast, 11 kidney and 11 pancreatic cell lines. For each cell line we selected all genes with a likelihood score higher than 0.65 in the Tsherniak study as a gene upon which its host cell is dependent, a total of 4,030 gene dependencies across 39 cell lines.

### Gene dependency count and magnitude of genomic alteration are significantly correlated

We first set out to find out if and how acquired gene dependencies differ across cell lines and tissue types and how gene dependency is related to genomic alteration. Using the data sourced via Tsherniak et al. we first plotted the number of gene dependencies reported for each cell line against a measure of that cell line’s genomic alteration.

We measured each cell line’s level of genomic alteration by counting the number of genes that had pathogenic mutations as identified by SIFT (Sim *et al*, 2012) and the number of genes differentially expressed when compared to the mean expression level for cell lines of that tissue type, using a cut-off point of 0.5 TPM.

Across all cell lines we found a slight but significant positive correlation between the measure of genetic alteration and the number of gene dependencies in cell lines (R= 0.36, p=0.012) (figure EV2 a.). To calculate the significance of this level of correlation we shuffled the data for genomic alterations 10,000 times, calculating the correlation coefficient each time to provide a normal distribution of correlation coefficients (figure EV 2. b.).

This significant positive correlation may be the result of alterations that have affected one or more otherwise non-essential genes that are part of synthetic lethal genetic interactions rendering the surviving gene in the pair as essential for cell viability.

We found that when compared to the other two tissue types cell lines originating in breast tissues exhibited, on average, a higher level of genomic alteration (p=3e-5) and a higher number of reported gene dependencies (p=0.024).

### Gene dependency signatures are enriched for specific disease tissue types

In order to quantify how gene dependencies are distributed across specific tissue types we next performed non-negative matrix factorisation (NMF) in order to find common signatures of gene dependency. To better understand how these signatures relate across tissue types we added additional cell lines from pancreatic tissue samples. To render the data more easily manageable for NMF we filtered our gene dependency data to remove genes that showed low variation between tissue types, i.e. any genes with var <0.1 across all tissue types were removed from the data before factorisation.

We found that 6 signatures was the minimum number required to describe the majority of the data (figure 1. a.). We took the most representational signature for each cell line and called this the cell line’s prominent signature. We plotted a count of cell lines with each corresponding prominent signature which was further grouped for tissue type to find enrichment. We found that two signatures contained only one type of tissue type, signature 2 which features only breast and signature 5 which features only kidney tissue. Signature 3 was also highly enriched for pancreatic tissue (figure 1. b.).

**Figure 1.**
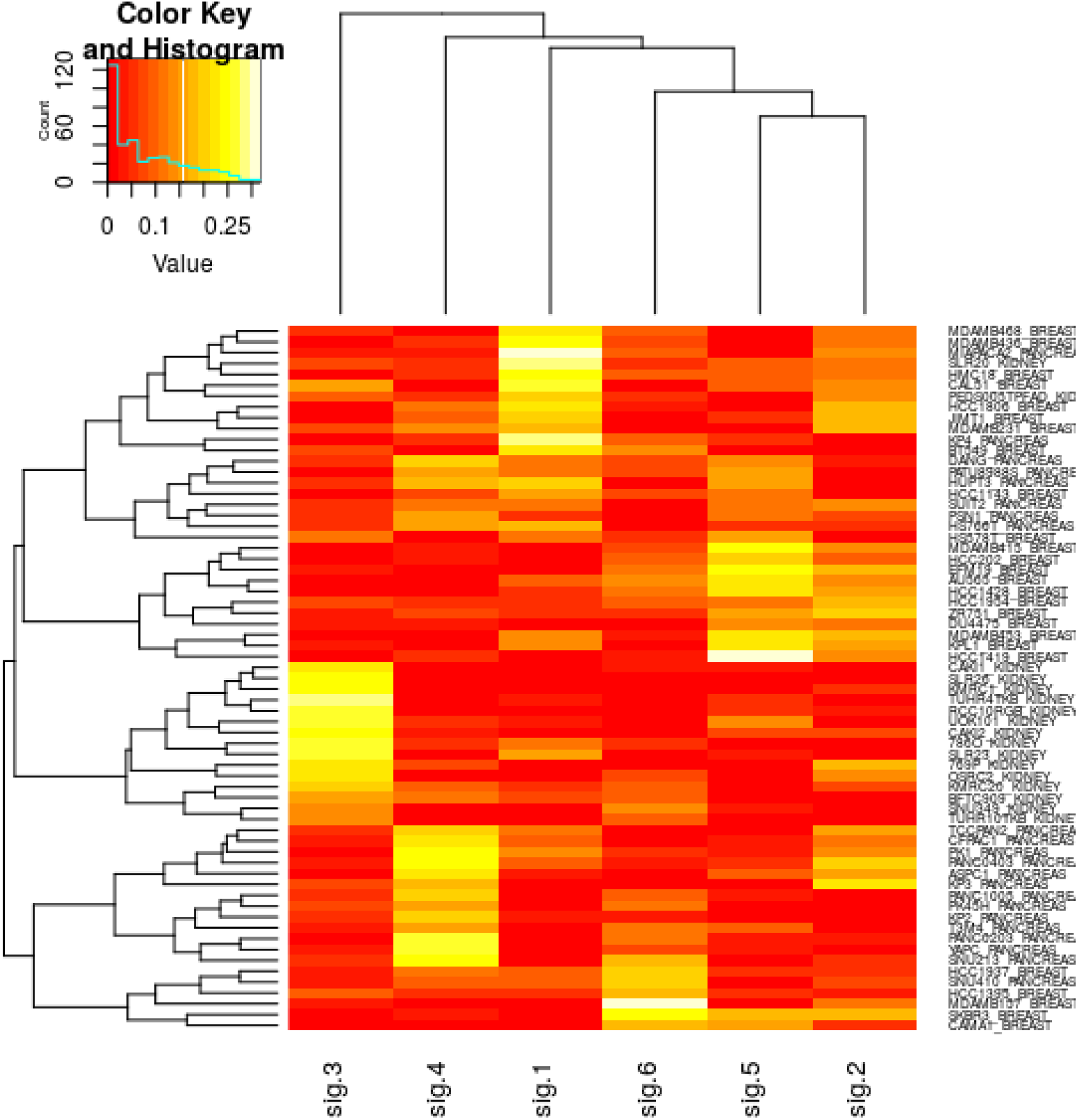
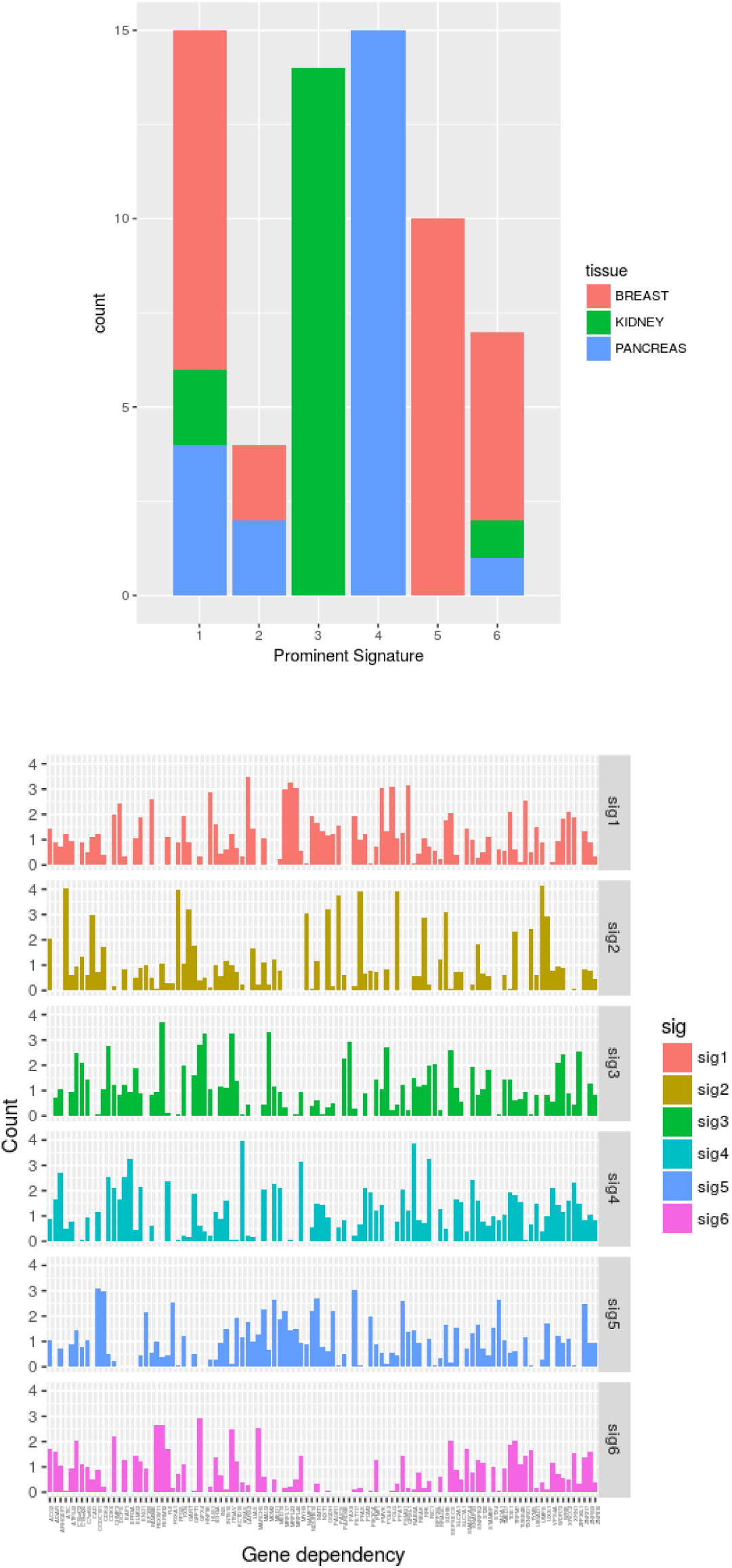
Genedependency signatures derived from non-negative matrix factorisation. a. A clustered heatmap shows the clustering of gene dependency signature prominence across cell lines. Dependency signature prominence sourced via the basis matrix (also known as matrix W) given by negative matrix factorisation. b. Enrichment analysis shows that tissue type is predictive of prominent gene dependency signature. Signature 6 for example is fully enriched for kidney cell lines, signature 2 for breast and signature 3 prominently features pancreatic cell lines. c. The composition of each gene dependency signature given by the mixture coefficients matrix (or matrix H)

This may suggest that different tissue types feature fairly stable, unique patterns of gene dependency either as a result of cellular environment or, especially in the case of cancer cell lines, synthetic lethal interactions.

For each signature we generated a list of the most prominently differentiated genes (figure 1. c.) by ranking the distance of each gene’s occurrence count in each signature from the from mean number of occurrences of that gene across all signatures as reported in table 1.

**Table 1.** Prominently differentiated genes between gene dependency signatures. For each signature every gene was ranked by distance from the mean score given by the basis matrix compared to the same gene across all other signatures.

| Sig 1 | Sig 2 | Sig 3 | Sig 4 | Sig 5 | Sig 6 |
| --- | --- | --- | --- | --- | --- |
| GART | FOXA1 | EFR3A | POLG | ELMO2 | PAX8 |
| ATIC | STX4 | KRAS | MRPL23 | GPX4 | MDM2 |
| CAD | MARCH5 | TUBB4B | MRPL46 | ITGAV | HNF1B |
| PAICS | TADA1 | RAB6A | HUS1 | VPS4A | RPP25L |
| PFAS | EP300 | SLC7A1 | LARS2 | FERMT2 | PARD6B |
| NAMPT | FBXW11 | MYH9 | MRPL17 | SEPSECS | ZFP36L1 |
| FPGS | CCDC101 | ARHGEF7 | QRSL1 | MARCH5 | POLE3 |
| UMPS | PIK3CA | VPS4A | DCPS | CHMP3 | CDK6 |
| LIAS | CDK4 | ADAR | PMVK | UBIAD1 | FERMT2 |
| OGDH | MED1 | EAF1 | TXNRD1 | SMARCA4 | C16orf72 |

### Modelling cell lines with biological network and genetic alteration data

For each selected cell line a model was created from the STRING PPI networks data (von Mering *et al*, 2005). In each model a node represents a protein and each edge between nodes a physical interaction between the two respective proteins. Once each model is generated in this way we essentially treat each node as the gene associated with the protein (figure 2.).

**Figure 2.**
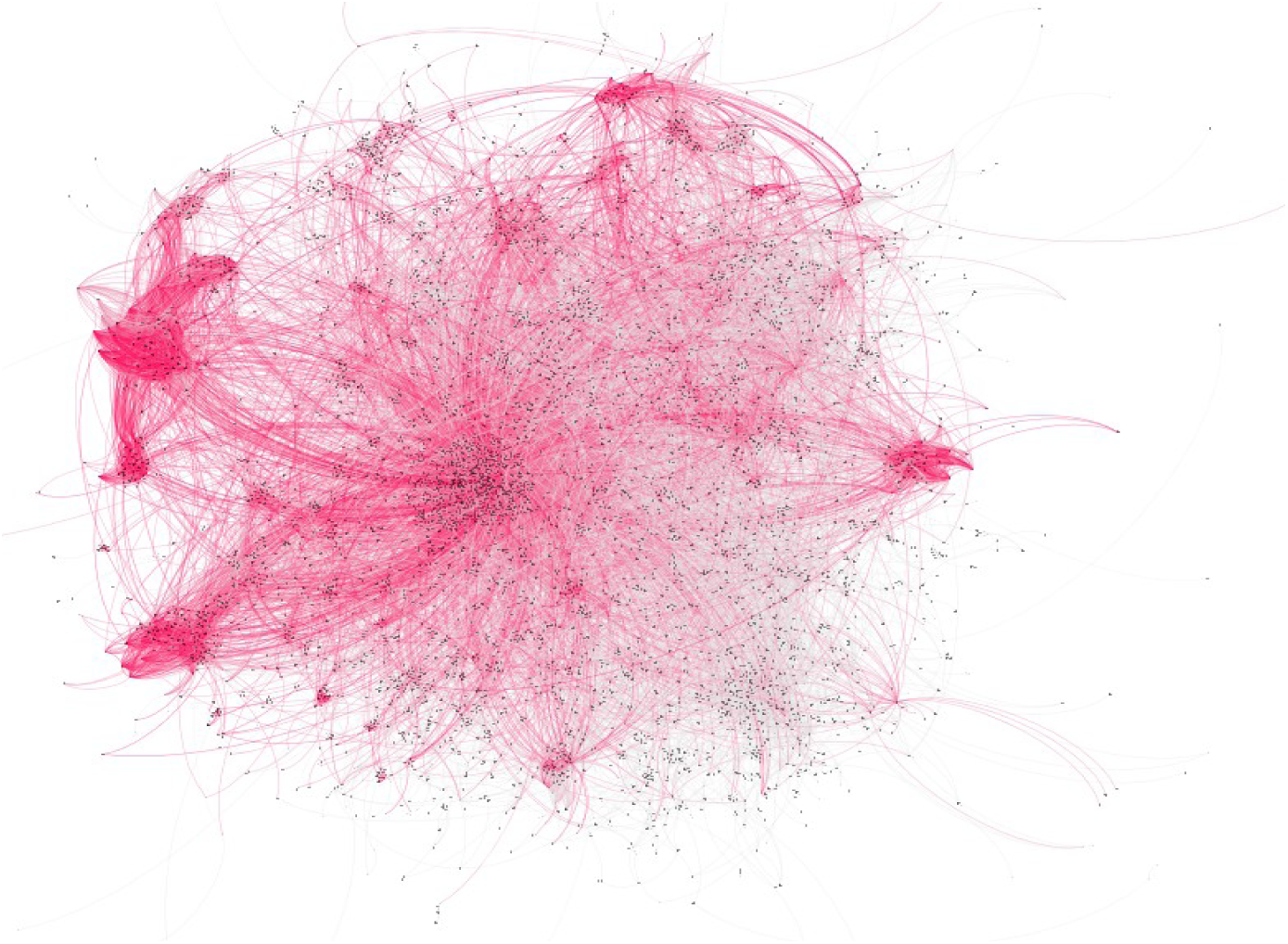
a. Plots of the PPI network graphs for breast cell line AU565 BREAST highlighting acquired essential genes in red suggests clustering of these gene dependencies.

We then extracted topology data for each node (table 2.) and used these data points as features in our machine learning models. The distribution of features values for dependency genes are somewhat different to those of non-dependency genes notably for the betweenness, constraint, eigen centrality and hub_score features (figure EV 3.) suggesting these features should provide some predictive power.

**Table 2.** List of graph topology features extracted from protein interaction network data with descriptions

| Feature name | Description |
| --- | --- |
| Betweenness | The number of shortest paths in the entire graph that pass through the node. |
| Constraint | Related to ego networks. A measure of how much a node's connections are focused on single cluster of neighbours. |
| Closeness | The number of steps required to reach all other nodes from a given node. |
| Coreness | Whether a node is part of the k-core of the full graph, the k-core being a maximal sub-graph in which each node has at least degree k. |
| Degree | The number of edges coming in to or out of the node. |
| Eccentricity | The shortest path distance from the node farthest from the given node. |
| Eigen centrality | A measure of how well connected a given node is to other well-connected nodes. |
| Hub score | Related to the concepts of hubs and authorities the hub score is a measure of how many well linked hubs the nodes is linked to. |
| Neighbourhood n size | The number of nodes within n steps of a given node for n of 1, 2, 5 and 6 |

We labelled the nodes in *PPI* network using the gene dependency data sourced via Tsherniak et al. (Tsherniak *et al*, 2017) for each cell line as either a dependency or non-dependency for training and validation. We refer to this unperturbed labelled PPI model as our base PPI networks model below.

### Base PPI network parameter data predicts pan-cell line dependency genes

To establish baseline performance for our classification models and to generate a list of relatively common dependency genes across cell lines we ran our classifiers on each cell line with no alterations or perturbations using the base PPI network discussed above.

We ran these classifiers to validate performance within cell lines, across cell lines of the same tissue type and across cell lines originating from different tissue types to understand how well the classifiers generalise.

To validate classification within individual cell lines we optimised our ADA boost classifiers’ hyper parameters using 5-fold cross-validation on our training data and further validated the classification performance using hold-out test data which constituted 20% of the full data set.

We validated the model on each of our 42 cell lines separately, using both training data and validation data extracted from the same single cell line. Each trial was repeated 10 times using the base PPI model. This gave us a mean predictive performance of AUC ROC 0.765 (s.d. 0.024).

To measure performance across cell lines originating from the same tissue type and the predictive performance between tissue types we used the training sets that were already generated for each cell line to train our classifiers and we systematically validated against each other cell lines test set.

To ensure that our models were not being biased by genes that were present in both training and test sets we ensured that any genes present in the training set were removed from the active test set.

We first measured how well our models generalise from one cell line to another within the same tissue type. Under these conditions the base PPI models had an average AUC ROC of 0.761 (s.d. 0.005), 0.755 (s.d. 0.008) and 0.754 (s.d. 0.012) for breast, kidney and pancreatic cell line sets respectively.

Finally, we trained our model on kidney data before predicting acquired gene dependencies in breast and pancreatic tissue. These cross cell line predictions resulted in a mean AUC ROC of 0.758 (s.d. 0.007) and 0.758 (s.d. 0.01) respectively. Similarly when we trained the model on breast data before predicting dependency genes in kidney tissue the model had a mean AUC ROC of 0.759 (s.d. 0.006) and breast to pancreas performed similarly with 0.761 (s.d. 0.01) Taking the mean performance of all cell lines predicting all other cell lines the base PPI network model gave an AUC ROC of 0.757 (s.d. 0.007).

### Feature importance

To quantify which features provide the most predictive power to our models we calculated a normalised importance score for each feature for each cell line and took the distribution of these scores across all cell lines. Feature importance was calculated by measuring the mean decrease in accuracy without each feature across all tree permutations in a random forest.

We found that a number of features that measure connectivity of a gene perform better than degree centrality although degree centrality does provide a moderate amount of predictive power. Page rank and eigen centrality scored well in all cell line models followed by hub score and constraint. Eccentricity, the distance a given node is away from the furthest node from itself in the network, a measure of how close that node is to the centre of the network, performs badly across all models.

These importance scores reflected the class feature distributions fairly well, i.e. features whose values varied more between essential and non-essential genes provided more predictive power.

Pagerank and constraint showed a noticeable differentiation between classes while the differentiation between classes for eigen centrality and hubscore features were not as prominent (figure EV 4.).

### Our perturbed models reported improved predictive power compared to our base model

Our base PPI models performed moderately well when predicting commonly observed essential genes within and across cancer cell lines. In order to improve overall performance and classify less common dependency genes that occur in a smaller subset of cell lines we used genetic alteration data to create unique models for each cell line.

Based on the available project Achilles mutation and expression data we applied a number of treatments to the base PPI networks to encode each cell line’s unique genetic alteration profile as discussed below.

Mutations such as frameshift indels or nonsense substitutions were labelled as loss of function. For missense mutations the Pathogenic mutations were identified using the SIFT online (Sim *et al*, 2012) and then split into either loss of function or gain of function using the MoKCaRF (Baeissa, 2019) algorithm. Nodes that represented genes with inactivating mutations were removed from the PPI network, for those that represented gain of function we amended the weights of their outgoing edges as discussed below.

As well as removing inactivated nodes we weighted edges to represent the strength of the signal between the two genes - the stronger the signal the lower the barrier. Two unidirectional edges were created between each gene pair (g1,g2).

We calculated each edge weight so that as gene expression (g) tends to 0, weight (w) tends to 1. As g tends to infinity, w tends to 0. Specifically

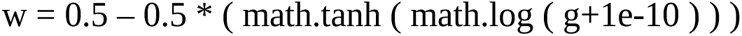

For genes subject to a gain of function mutation we multiplied the gene expression by 10 before calculating the weight. Whilst the exact equation w is somewhat arbitrary we found that our results were robust to changes in w.

We used three distinct versions of our expression data to accomplish these perturbations. We first used the raw expression data for each gene directly, next we normalised the expression level of each gene in a cell line against the same gene in all other cell lines of the same tissue type and finally, we normalised the data against the same gene in all other cell lines.

We found that of all the PPI networks treatments the raw gene expression data showed the best overall predictive performance both within and across cell lines. Within cell lines our raw data models scored a mean AUC ROC of 0.812 (s.d. 0.023) compared to the base model’s performance of AUC ROC 0.765 (s.d. 0.024).

Predicting across all cell lines and all rarities of gene our raw data model performed with ROC AUC of 0.801 (s.d. 0.006) again an improvement performance to that of the base PPI networks model’s mean ROC AUC of 0.758 (s.d. 0.007) (figure 3.) (Table 3.).

**Figure 3.**
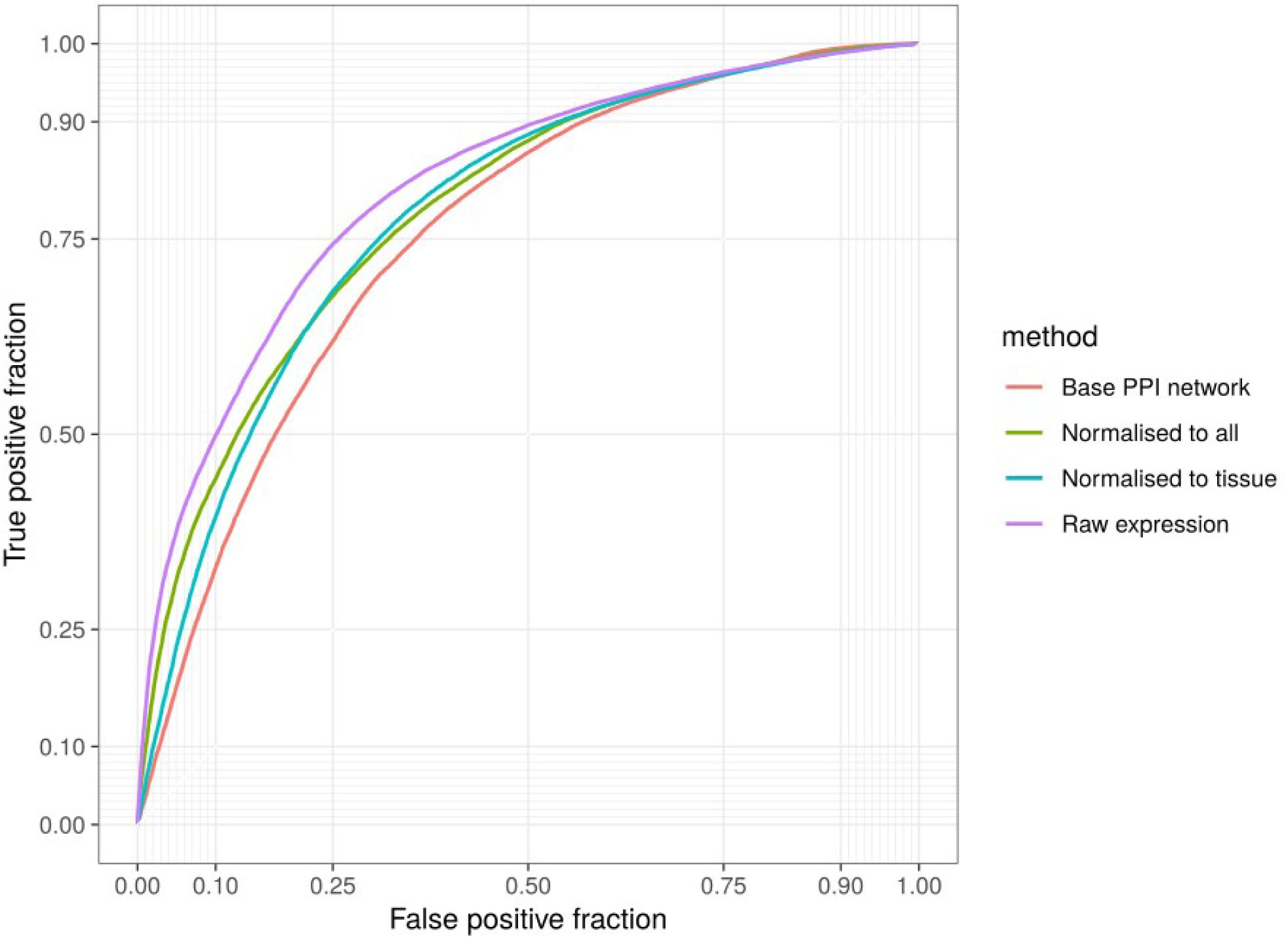
AUC ROC plots for each PPI model show that our raw expression model exhibits the largest AUC ROC, and therefor the best performance, while the base PPI model shows the worst performance.

**Table 3.** Mean model performance when predicting gene dependencies within each cell line (where training and test datasets were sourced from a single cell line) and across cell lines (where training was sourced from one cell line and used to classify all other cell lines). Performance measured with mean AUC ROC scores.

| **PPI networks treatment** | **Description** | **Within Cell line mean AUC ROC** | **Across all cell lines mean AUC ROC** |
| --- | --- | --- | --- |
| Base PPI network | Non-directional PPI network sourced via STRING. | 0.765 (s.d. 0.024) | 0.758 (s.d. 0.007). |
| Compared to tissue type | Gene expression normalised against all cell lines of same tissue type. | 0.778 (s.d. 0.021) | 0.756 (s.d. 0.128) |
| Compared to all | Gene expression normalised against all cell lines. | 0.781(s.d. 0.023) | 0.756 (s.d. 0.01) |
| Raw data | Raw gene expression data via Cancer cell line encyclopaedia (CCLE). | 0.812 (s.d. 0.023) | 0.801 (s.d. 0.006) |
| **PPI networks treatment** | **Description** | **Within Cell line mean AUC ROC** | **Across all cell lines mean AUC ROC** |
| Base PPI network | Non-directional PPI network sourced via STRING. | 0.765 (s.d. 0.024) | 0.758 (s.d. 0.007). |
| Compared to tissue type | Gene expression normalised against all cell lines of same tissue type. | 0.778 (s.d. 0.021) | 0.756 (s.d. 0.128) |
| Compared to all | Gene expression normalised against all cell lines. | 0.781(s.d. 0.023) | 0.756 (s.d. 0.01) |
| Raw data | Raw gene expression data via Cancer cell line encyclopaedia (CCLE). | 0.812 (s.d. 0.023) | 0.801 (s.d. 0.006) |

### Perturbed PPI network models perform well for both common and rarer gene dependencies across cell lines

To quantify how well our models predict those genes with high dependency scores in only a few cell lines we trained our models on all cell lines and then performed validation on test sets filtered for the rarity of the acquired essential genes being predicted. 580 of the total 4030 (∼14.3%) essential genes in our training data were identified as essential in all 39 cell lines. 2424 (∼60.1%) were essential in more than half of the cell lines and 821 (∼20.3%) of the total genes were specific to just one cell line. We created test sets featuring genes that occurred in just 1 cell line, below 10, 20, 30 in all 39 cell lines to calculate how well our models performed at each gene dependency rarity interval.

Of our four models, three (our base PPI network, proportional to tissue and proportional to all models) had similar levels of predictive ability for gene dependencies found in all cell lines in our training data. Across the other rarity intervals though the proportional models performed slightly better than the base PPI model.

The final model, our raw expression model outperformed the other models by some margin reporting a mean ROC AUC 0.660 when predicting genes that were reported as a dependency in only 1 cell line (compared to base model’s 0.615), 0.681 for genes that showed dependency in less than 10 cell lines (compared to 0.621), 0.711 (compared to 0.644) for genes in less than 20 cell lines, 0.727 (compared to 0.665) for genes in less than 30 cell lines and 0.801 for all gene dependency rarities (compared to 0.758 for the base PPI model) (figure 4.).

**Figure 4.**
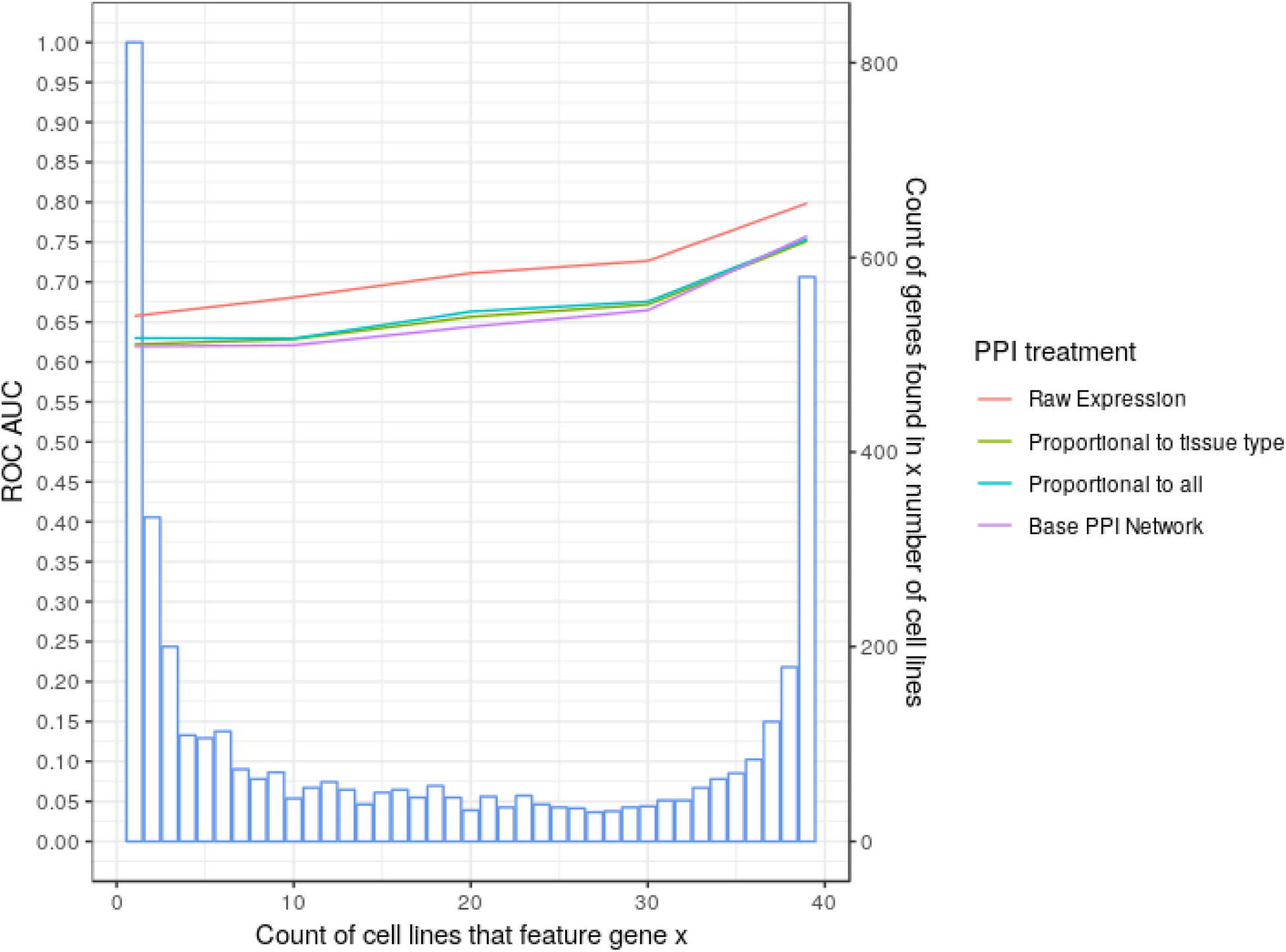
Model performance across gene dependency rarity intervals shows the general improved perforamnce of the raw expression model. Each coloured line represent a models performance at each interval as per the legend and the blue bars represent the distribution of genes at each rarity level. For example 200 genes are reported to be dependency genes in exactly 3 cell lines.

### Our models are fairly robustness to PPI networks incompleteness

It is known that current PPI networks models are both incomplete and suffer from ascertainment bias in that some proteins are better studied than others (Mosca *et al*, 2013; Rolland *et al*, 2014; Huttlin *et al*, 2017). In order to quantify how the incomplete nature of the PPI networks affects the robustness of our models, we repeated our classification pipelines with revised PPI networks data randomly holding out 25% of the data from original network. In the case of the 25% holdout PPI networks network we observed minimal loss of predictive power from our raw expression cross cell line model with mean reported performances of AUC ROC 0.78 (s.d. 0.011) from and 0.801 (s.d. 0.006).

We conclude that while an increasingly complete PPI network may improve their predictive performance our current models are fairly resilient to the incomplete nature of the currently available PPI networks data.

### Creating a pan tissue cell line training set

To maximize the amount of training data available for use by our classifiers for the prediction of gene dependencies in previously unlabelled cell lines we concatenated all available cell line training sets from all tissue types into one super set. We used our raw expression models for this super set based on their relatively high overall performance during previous validation.

In an attempt to estimate how well this concatenated data should perform for the prediction of gene dependency in unlabelled data sets we once more validated each of our individual test sets based on models trained using our super training set.

We found that our super training set classified gene dependencies across all cell lines with an AUC ROC of 0.843 (s.d. 0.012), a further improvement on the individual raw expression models mean cross cell-line AUC ROC score of 0.801 (s.d. 0.006).

This model provided the most predictive power and as such represents the most suitable available for predicting gene dependencies in cell lines with no prior labeling as discussed below.

### Predicting and validating gene dependencies in previously unlabelled cancer cell lines

To create a set of predictions we took 38 cell lines previously unlabelled for gene dependency, 16 for breast, 13 for kidney and 8 for pancreas. Each of these cell lines were chosen based on the amount of mutation and expression training data available. We used our pan-tissue training set to train our classifiers and produced a full set of predictions for each of these cell lines.

Survival screens focusing on a library of 240 genes involved in the DNA damage response (DDR) were repeated in triplicate for the MCF7 breast cell line. Cell viability was reported using a z-score where positive numbers suggested a cell’s viability increases with the knockdown of the predicted gene, negative scores suggests a decrease in viability and z-scores below -1 constitute a true dependency. The variance of results across all three repeats was high. This may have been due to the choice of library. The loss of genes involved in the DDR can often lead to genomic instability in a cell. Knocking out a single gene (eg MSH3) can cause the subsequent loss of different sets of genes, resulting in different sets of dependencies.

We ranked all of our predictions for MCF7 by dependency likelihood score. Filtering for likelihood scores to keep predictions of above 0.85 and below 0.15 and treating negative z-scores as a hit we report an accuracy of 0.64 with a sensitivity of 0.73 and a false discovery rate of 0.38 based on experimental validation for the MCF7 cell line. Next, we extracted the top 10 predictions. 8 of our top 10 predictions showed signs of essentially with a mean negative z-score. Two of these top 10 predictions, PARP1 and TRIM28, reported a z-score of less than -1 in at least one repeat (table 4). Only 7 of the 240 genes screened and classified for in the MCF7 cell line reported a mean z-score of less than -1 in all 3 repeats and 2 of these, MEN1 and CHEK1 were predicted as gene dependencies with a score of over 0.85 (table 5).

**Table 4.**
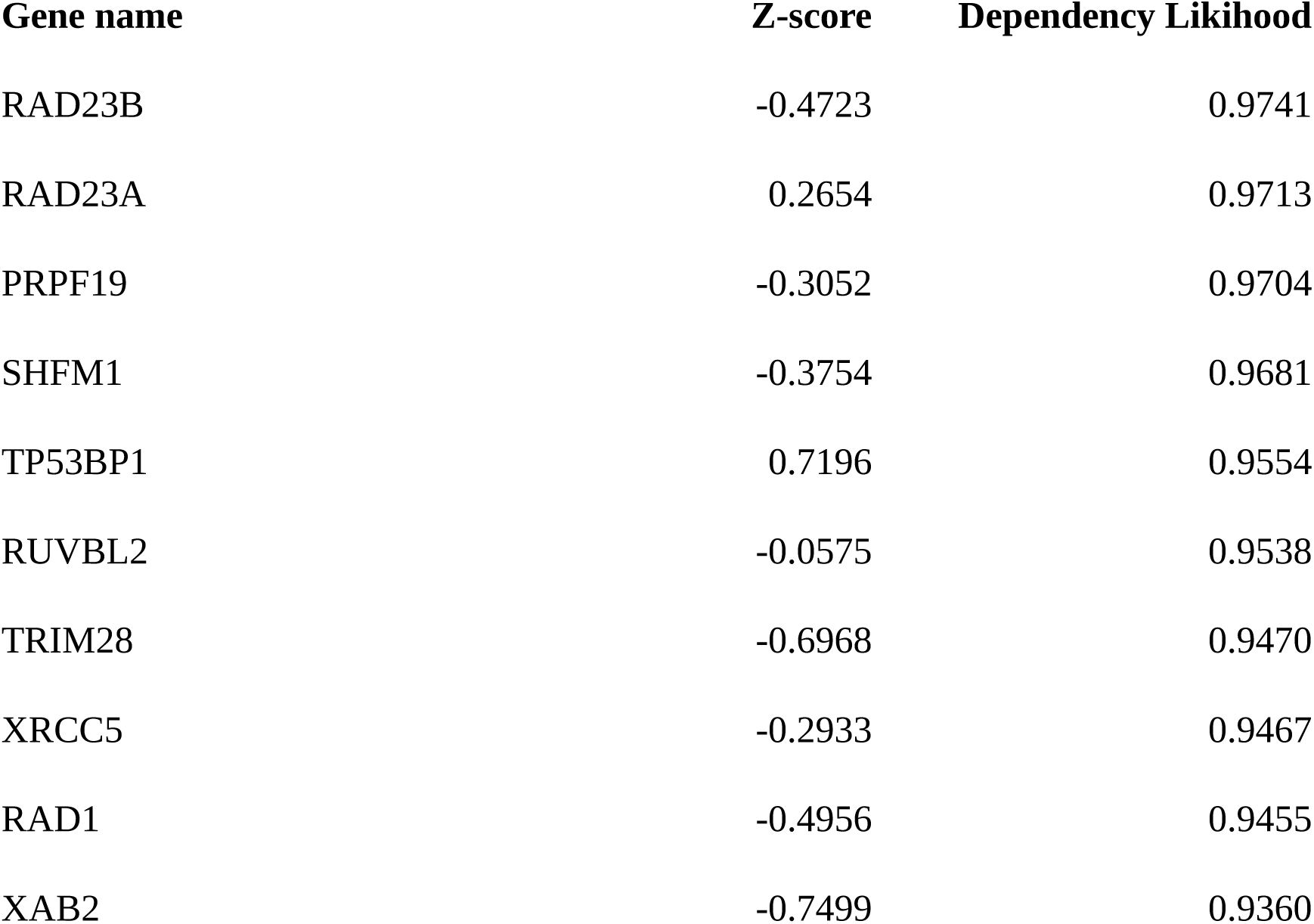
Top 10 dependency gene predictions with likelihood score reported by our pan-tissue classifier and z-scores from the MCF7 DDR library survival screens. Negative z-scores suggest that the knockout of a predicted gene impacts cell viability and z-scores of below 1 suggest dependency. 8 of these 10 genes showed negative z-score with XAB and TRIM28 reporting a z-score of less than -1 in at least one repeat of the screen.

**Table 5.**
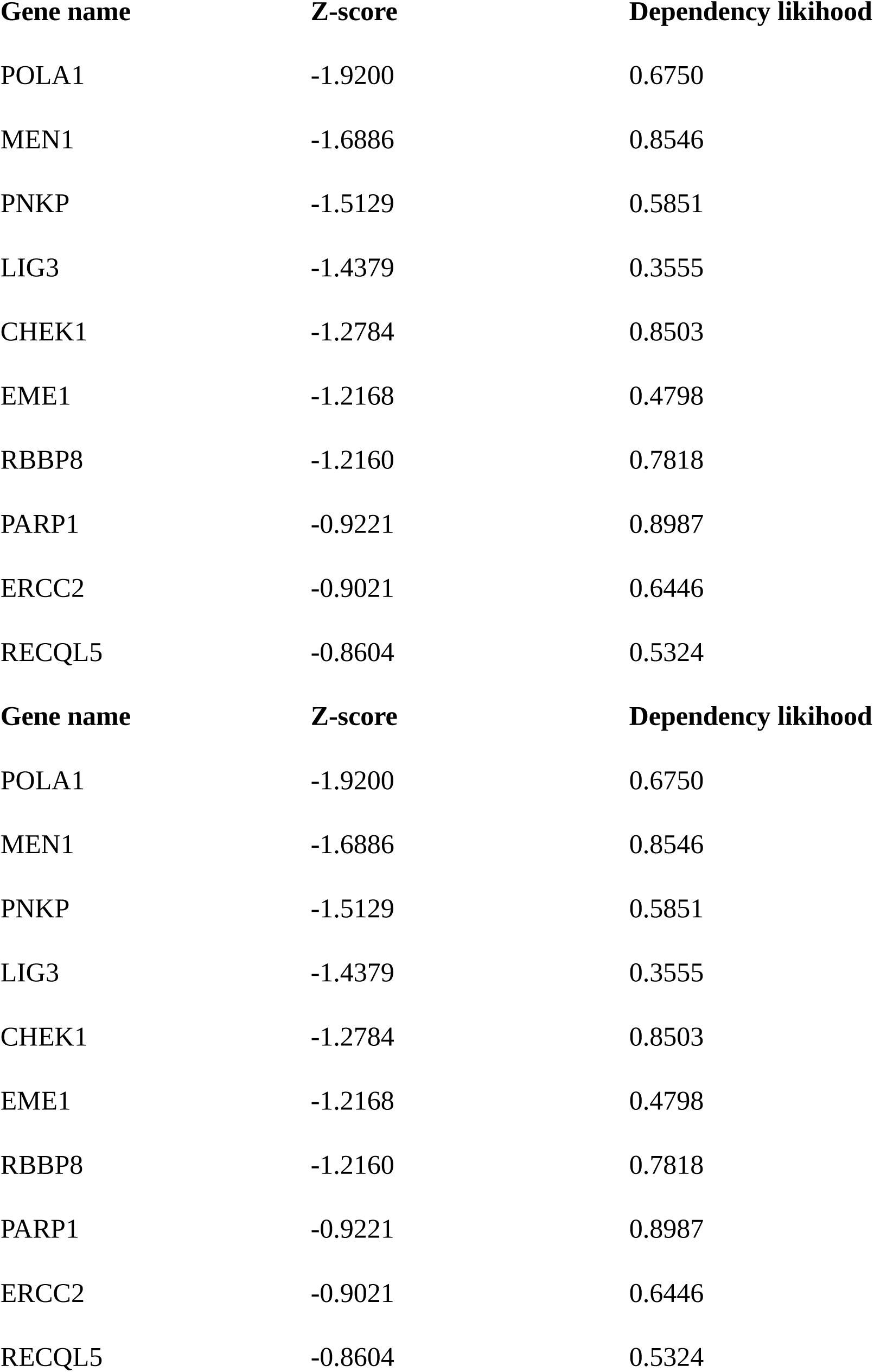
The 10 lowest genes by reported z-score in the MCF7 cell line with dependency likelihood scores given by our pan-cancer classifier. 3 of these, MEN1, CHEK1 and PARP1 obtained dependency likelihoods of over 0.85 and 8 of the 10 scored over 0.5.

### Therapuetic opportunites in cancer dependency genes

Using Cansar’s cancer protein annotation tools (Bulusu *et al*, 2014) we labelled our predicted dependency genes, based on their respective protein products, as either a drug target, druggable or non-druggable (Figure 5). The proportion of known drug targets in our predicted gene set was slightly lower than those in our training data at 0.7% compared to 1.1%. The proportion of predicted druggable genes based on three dimentional structure was higher at 45.1% compared to 34.2% in our training set. We found therapuetic opportunities in almost every cell-line in both our training data and prediction set both in the form of genes with known drugs and genes that exhibit druggable traits.

**Figure 5.**
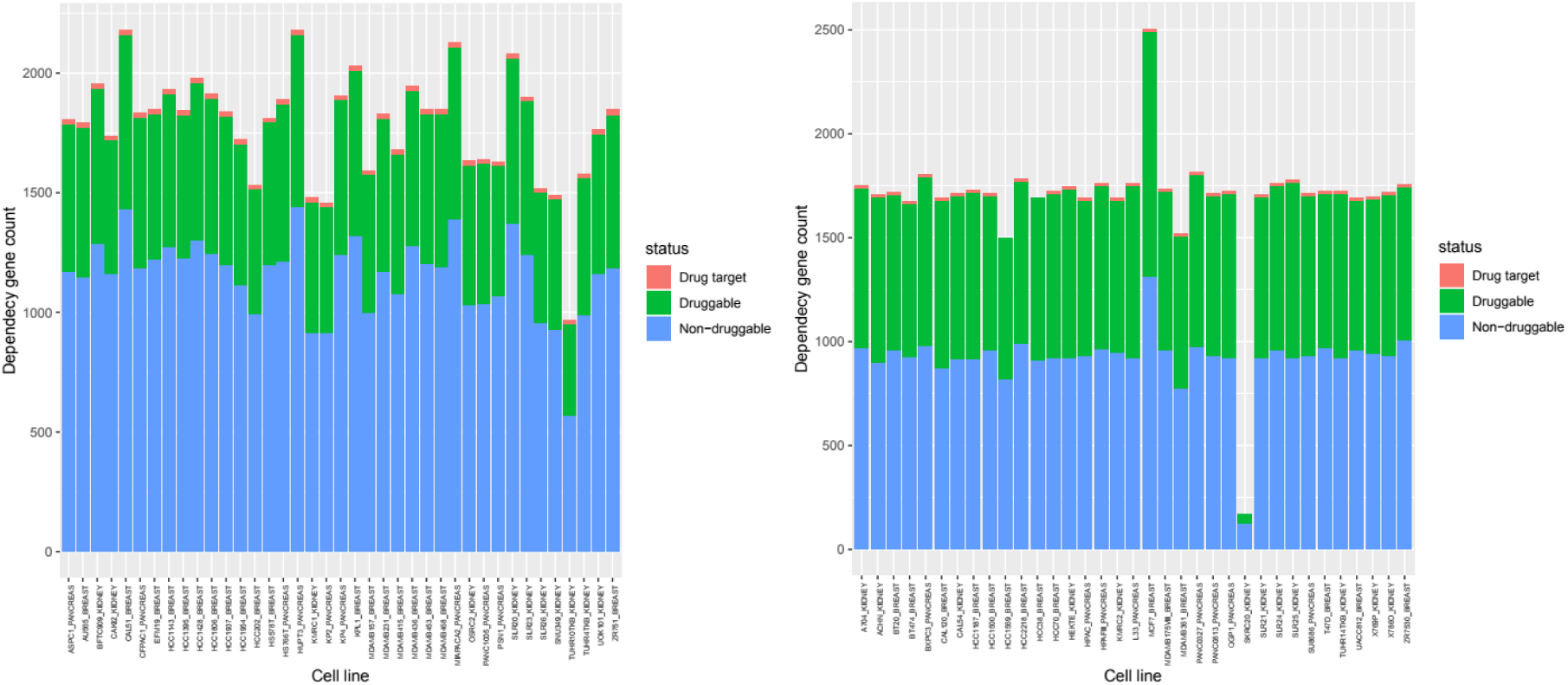
Dependency gene druggability counts by cell line. a. A histogram of dependency gene counts per cell line in our training data stratified by druggability status as reported by cansar black’s cancer protein annotation tools. b. Predicted dependency gene druggability count by cell line.

## Discussion

Protein-protein interaction maps provide us with a robust model of how the proteome is organised. Here we find that the topological relationships across these maps tends to be different for essential genes and non-essential genes, opening up the opportunity for predicting gene dependency. We find that topological features can be used to predict gene dependency in human cell lines with ROC AUC scores of up to 0.84. This is an improvement on accuracy reported by previous studies that use PPI network models to predict essential genes in *S. cerevisiae* (Saha & Heber, 2006) and *E coli (da Silva *et al*, 2008)*.

Jeong et al’s seminal publication (Jeong *et al*, 2001) was the first to show a correlation between degree centrality, i.e. the number of edges leading in or out of a given node, and gene essentiality. We find here that it is possible to use these and other topological features to predict essential genes and acquired essential genes in previously unseen cell lines, using models trained on different cell lines. We note though that the topological features that are predictive of gene dependency such as eigencentrality are predominantly measures of a protein’s connectedness. These features are robust to the type of network perturbations caused by changes in gene expression and mutations. This suggests that modified PPI networks can only provide a partial picture of gene essentiality.

We described how the standard Protein Protein Interaction Network does not capture the massive cell reorganisation seen in cancer, due to genetic mutations, copy number variances and epigenetic changes affecting gene expression. By personalising our PPI networks to reflect some of these changes we were able to model our cells lines better and improve predictive power gene dependency classification. This improvement is particularly noticeable for those genes we are particularly interested in, ie the genes which are essential in only a few cell lines.

Despite the relatively high performance of our classifiers we are aware that the association between gene expression and protein expression is only partial and so it is likely that further improvements will be possible for this type of model when it is possible to modify the PPI network as a result of protein expression as well as existing ‘omic data.

Additionally consideration of the biological nature of the protein interactions reported as well as improvements to the completeness of our source PPI networks is also likely to lead to significant improvements in this type of study. In particular our source protein-protein interaction network provides only non-directional, binary information about interactions between proteins rather than the inhibitory or excitatory nature of the interaction. Although we report that our models are relatively robust to incompleteness in the source networks we expect that as the completeness and sophistication of PPI models improves so will the effectiveness of this type of model.

## Methods

### Constructing the base PPI

Our base protein-protein interaction data was obtained via the STRING database (v.10) (von Mering *et al*, 2005). This data was filtered to include only interactions with an experimental score higher that 80 to ensure each interaction was reliable. The ENSP protein IDs in this data set were converted to their respective ENSG gene IDs using Ensembl data (Hubbard *et al*, 2002). R (version 3.4.0) and the igraph package (version 1.1.2) (Csárdi & Nepusz, 2006) were used to produce a network model of the PPI data for each cell line.

### Essentiality data and labelling

The Tsherniak et al. (Tsherniak *et al*, 2017) survival screen data, via project Achilles, provides a likelihood score for each gene in each cell line being a essentiality. We the same likelihood threshold as Tsherniak et al. to label each gene in our model as a gene dependency, those above 0.65 or non-dependency those below 0.65 for each cell line.

### Perturbing the PPI

All edges in the directed PPI network have a weight in (0,1] which reflects the strength of expression of the initial protein, i.e. proteins that are not expressed have edges of weight 1 emanating from them, and as expression increases so the weight reduces. These weights are determined by modifying RNA seq data to reflect the loss and gain of function of proteins with mutated gene sequences.

In order to create these weights, RNA seq data from was downloaded from the Cancer Cell Line Encyclopaedia (Cancer & Line, 2015), and mutation data was downloaded from Tsherniak via Achilles (Tsherniak *et al*, 2017).

Mutations that lead to loss and gain of function were identified as follows. Frameshift indels were assumed to lead to loss of function. The program SIFT (Kumar *et al*, 2009) was then used to separate out the missense mutations that have no functional impact. The remaining missense mutations were categorised as leading to either loss of function or gain of function using a version of the MOKCARF algorithm. MOKCARF uses features from Mutation Assessor (Reva, B.A., Antipin, Y.A. and Sander, 2010), Polyphen2 (Adzhubei *et al*, 2013) and FATHMM (Shihab *et al*, 2013) as input to a ADA boost classifier which has been trained on protein domains mutated in proto-oncogenes, or tumour suppressor to identify loss or gain of function.

Gain of function is assumed to have a multiplicative impact on RNA expression (set here to a factor of 10), whilst loss of function sets the resulting weight to 1.

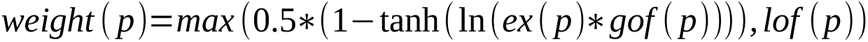

where ex(p) is the RNA_seq expression associated with protein p

*gof* (*p*)=10 if there is a mutation in the gene associated with protein p leading to gain of function, otherwise 1.

and *lof* (*p*)=1 if there is a mutation in the gene associated with protein p leading to loss of function, otherwise 0.

### Feature generation

R and the igraph package were used to extract 14 network topology features for each cell line’s protein interaction network described in table x.

### Preprocessing feature data

To improve performance in cross cell line classification each cell line’s feature set was normalised (Jacunski *et al*, 2015). To ensure unbiased validation we held-out 20% of this data to be used as a test set leaving 80% to be used as training data.

### Model validation

Classification was performed using the R caret library’s “ADA” boosted classification trees classifier. 5-fold cross validation was applied to each celllines training data to select the most optimised set of hyper-parameters. The ADA classifier as implemented in the caret library has three hyper-parameters to optimise, number of trees, max tree depth and learning rate.

A final model using these optimised hyper-parameters was then used to predict against the hold-out test set to assess predictive performance within each cell line and between each cell lines. These predictions were outputted as the probability of each class, essential or non-essential.

### Pan-cancer model and unlabelled predictions

To predict dependency genes in unlabelled cell lines we first concatenated all training data into one large labelled training data set. We produced a number of feature sets for cell lines that were not included in the original training data and predicted dependency genes in these unlabelled cell lines based on a model trained on the pan cancer set.

### Experimental validation

We chose a single unlabelled cell line, MCF7, for experimental validation. MCF7 was not featured in our training data and was chosen based on ready availability and good class balance for predictions on genes featured as part of the available DDR gene library.

We performed a high-throughput siRNA screen for experimental validation. Human breast (adenocarcinoma) MCF7 cells (validated by ATCC STR.V profiling) were grown in MEM supplemented with 10% FCS, penicillin/streptomycin and L-glutamine at 37oC and 5% CO2. Cells were reverse transfected with library siRNA using lipofectamine RNAiMAX (as per the manufacturer’s instructions) in black 96 well plates. Plates were incubated at 37°C, 5% CO2 for 72 hours. CellTitre-Blue was added to determine cell viability, plates were analysed using a plate reader at 560/590nm.

### Druggability annotation

Druggability annotation was performed using Cansar Black’s cancer protein annotation tools (Bulusu *et al*, 2014). We designated any genes with a “nearest drug target” score of 100% as a known drug target and any gene with one or more predicted drug targets in three dimentional structures that exhibited 100% homology with the respective gene’s sequence Identity.

## Acknowledgements

This work was supported Medical Research Council studentship [grant number MR/N50189X/1] (to G.B.-H) and funding from The Wellcome Trust [Institutional Strategic Support Fund 204833/Z/16/Z] (to A.M.C).

## EV Figures

**Figure EV 1.**
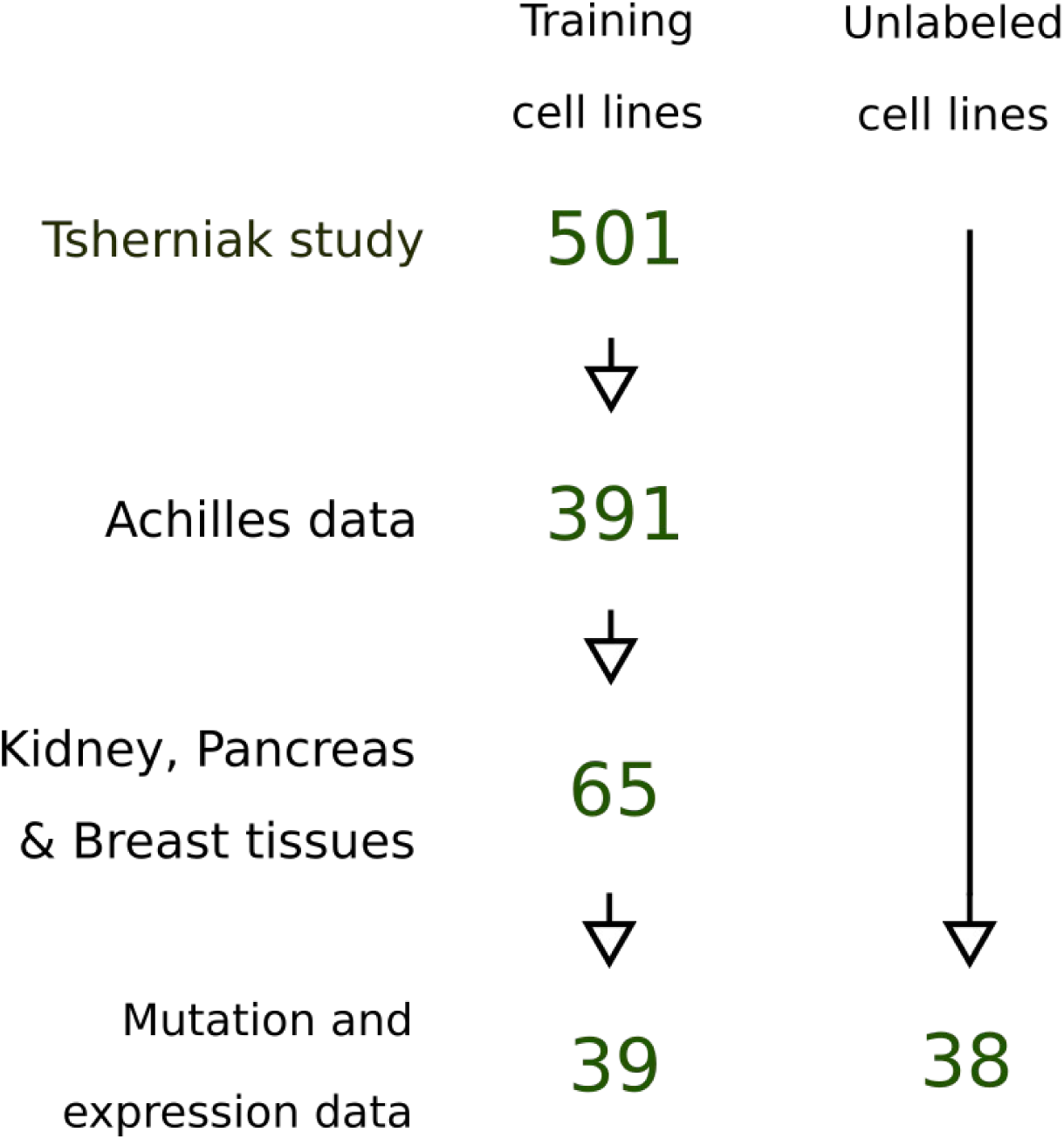
Number of cell lines available after filtering for public accessibility, tissue type and genetic alteration data availability.

**Figure EV 2.**
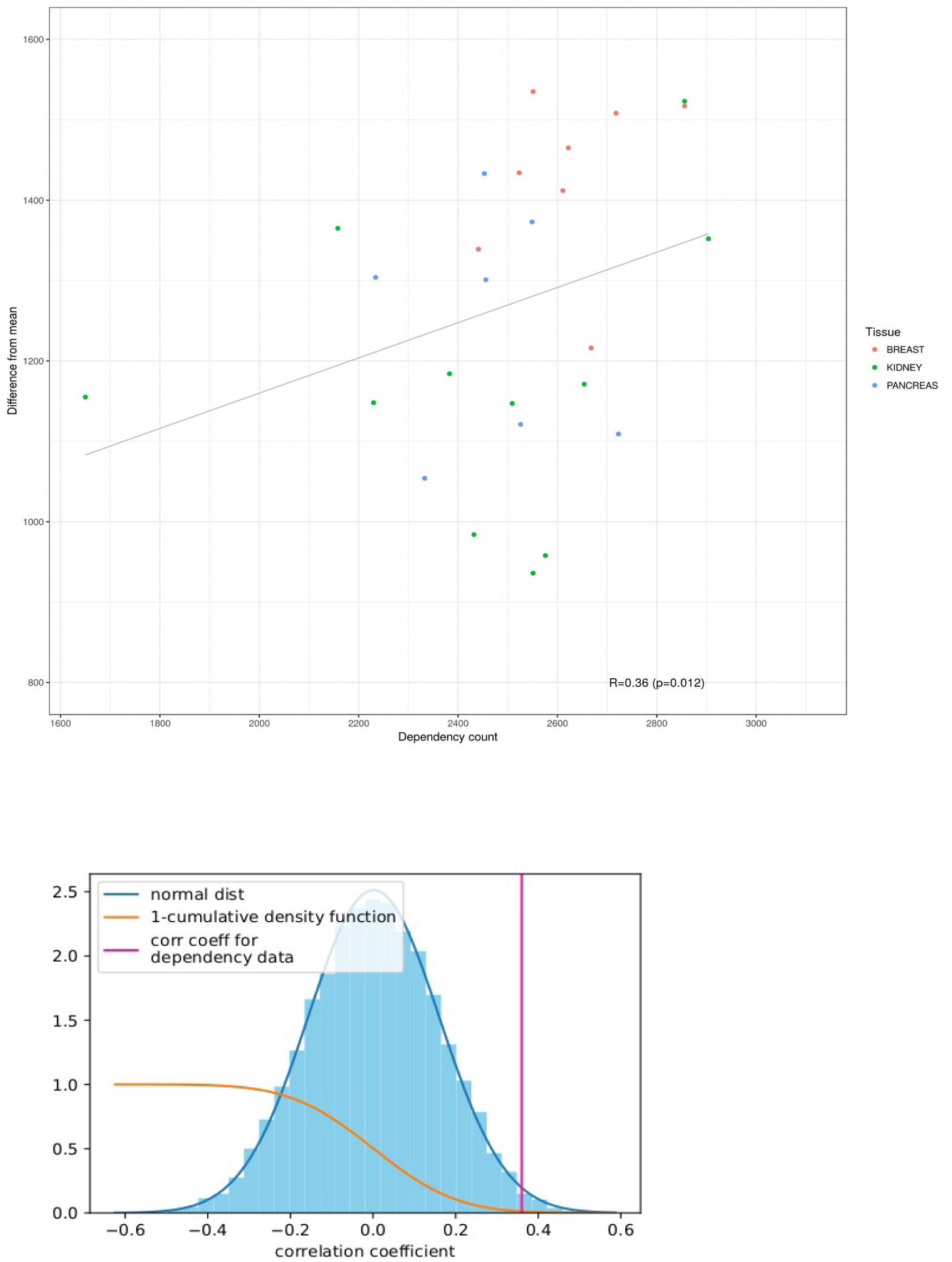
Measuring the relationship between generic alteration levels and count of gene dependencies in cell lines. a. By plotting the number of gene dependencies reported for each cell line against a measure of that cell lines genomic alteration we find a small positive correlation between the two. b. Shuffling the data and then finding the correlations for this data demonstrates that our correlation is statistically significant p-value=0.012.

**Figure EV 3.**
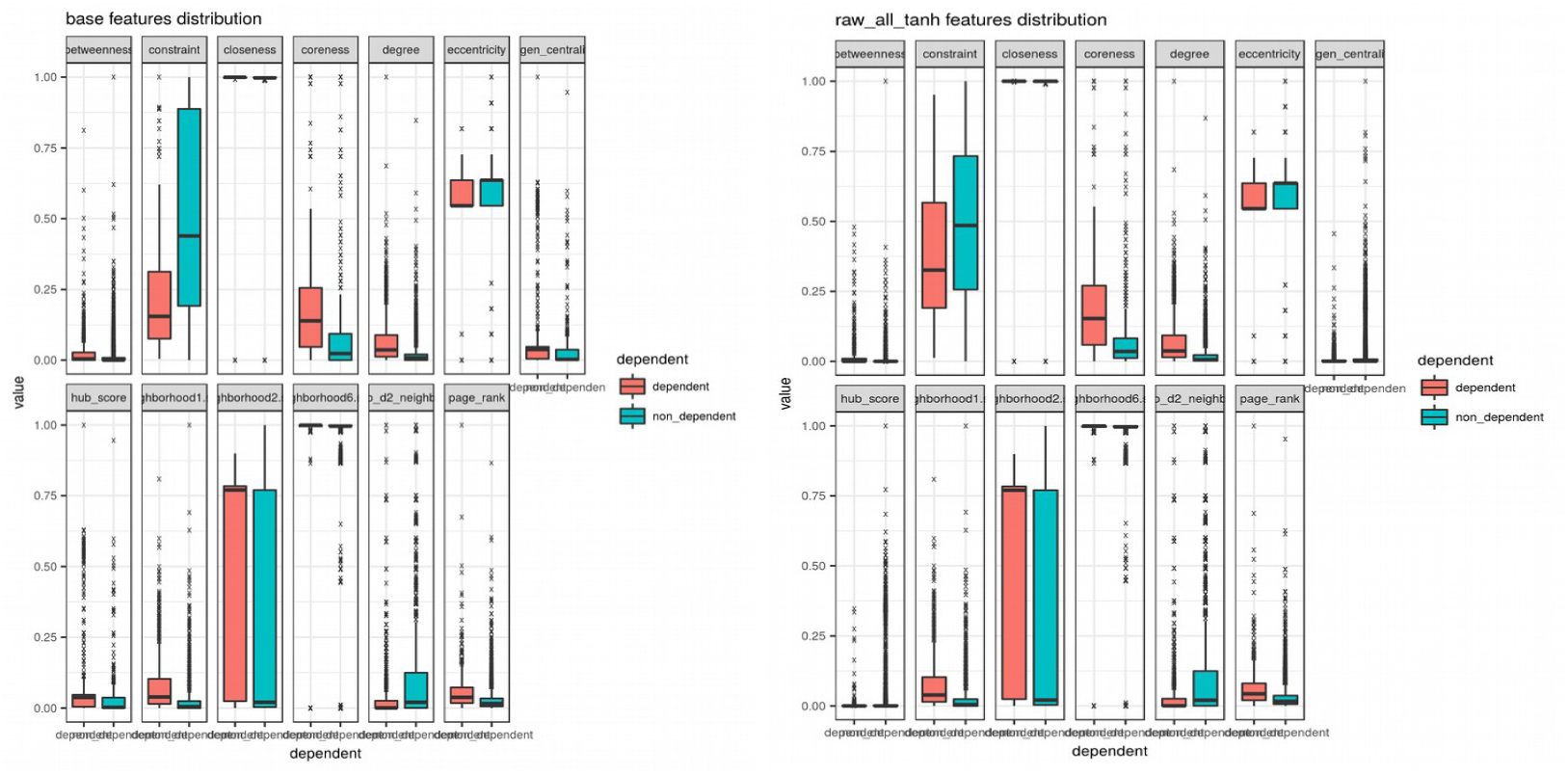
Feature distributions between dependent and non-dependent gene classes show some differences between the classes for the betweeness, constraint, eigen centrality and hub_score features

**Figure EV 4.**
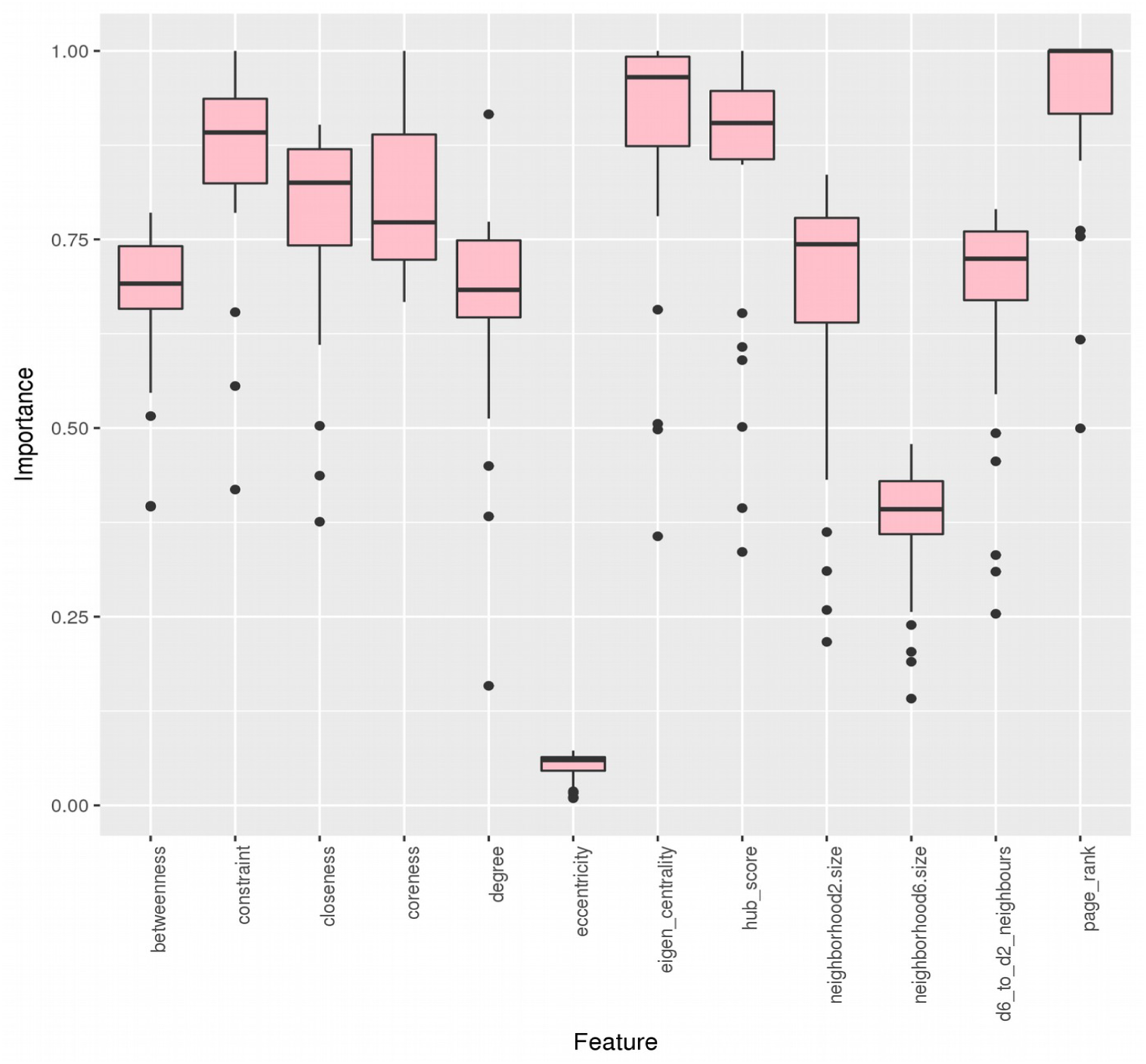
Importance for each feature used in each model calculated by measuring the mean decrease in accuracy when holding out each variable across all tree permutations in the random forest.

**Figure EV 5.**
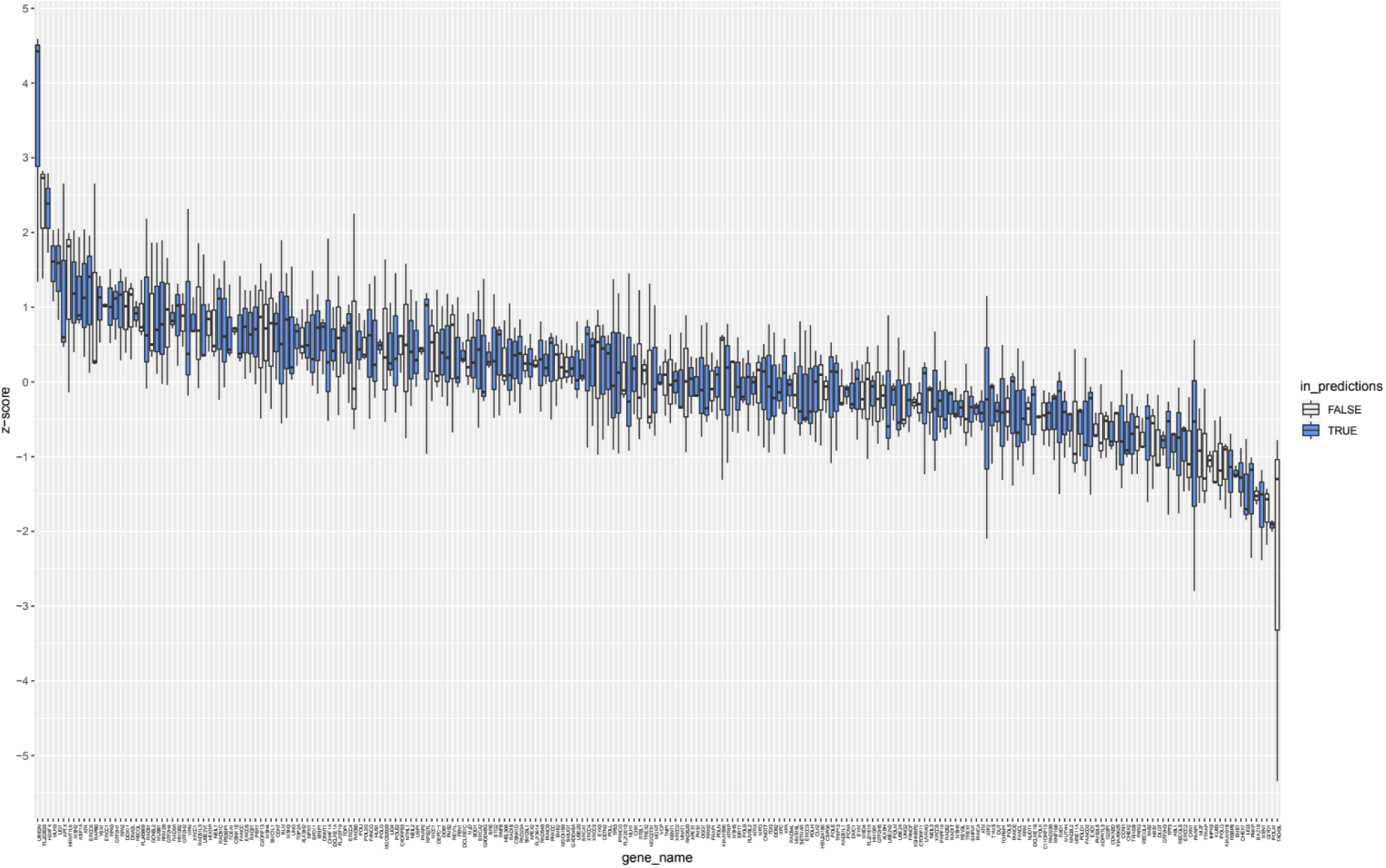
Survival screen’s z-score distribution with variation. This box plot graphs each gene featured in the survival screen with its z-score distribution across 3 experimental repeats. Blue boxes denote genes which were featured in our prediction set. White boxes denote genes that were not in our prediction set due to insufficient training data (i.e. mutational or copy number data).

